# Susceptibility of small grain cereals to *T. afroharzianum* ear infection in the field related to fungicide treatment and crop cultivar

**DOI:** 10.1101/2025.10.06.680620

**Authors:** Annette Pfordt, Susanne Pruszynski-Beck, Andreas von Tiedemann

## Abstract

Members of the genus *Trichoderma* are widely recognized as beneficial fungi and frequently applied as biocontrol agents. However, recent studies have identified *Trichoderma afroharzianum* as a pathogenic species, capable of infecting maize, wheat, and barley under greenhouse conditions. Its pathogenic potential in small-grain cereals under field conditions, however, has been so far unexplored. To evaluate its pathogenic potential, two-year field trials (2023 and 2024) were conducted using ten wheat, five barley, two rye, and one triticale cultivar. Artificial inoculation was carried out at full flowering using a spray inoculation with pathogenic *T. afroharzianum* isolates obtained from maize and water as control. Three fungicides were applied: untreated control, foliar treatment before inoculation, and ear application after inoculation. Disease development was assessed based on colonization rate of harvested kernels, and thousand kernel weight (TKW). The results showed significant differences in susceptibility between and within cereal species. *T. afroharzianum* was able to colonize wheat and barley ears under field conditions, with some cultivars showing significant reduction in TKW and high colonization rate. In contrast, rye and triticale exhibited much lower infection rates. Fungicide treatments had varying levels of efficacy. Ear application after flowering was most effective in reducing colonization and preventing yield loss, whereas early foliar treatment showed limited effect. These findings indicate that *T. afroharzianum* may represent a relevant pathogen in cereal production adding a new ear disease in cereals and requiring monitoring and enhanced fungicide applications in the future, unless less susceptible cultivars are identified and utilized in practice.

## 1. Introduction

Cereals such as wheat (*Triticum aestivum*), barley (*Hordeum vulgare*), rye (*Secale cereale*), and triticale (× *Triticosecale*) are among the most important staple crops worldwide, providing essential food and feed resources (FAOSTAT, 2021). Wheat is the most widely cultivated cereal globally, serving as a primary source of carbohydrates and protein for human consumption. Barley plays a crucial role in animal feed and the brewing industry, while rye is primarily used for bread and baked goods, it also serves as livestock feed and a key ingredient in whiskey and other alcoholic beverage (Wrigley et al., 2017). Triticale, a hybrid of wheat and rye, combines the beneficial agronomic traits of both parent species, offering improved resistance to environmental stress and better grain quality under suboptimal growing conditions (Eudes, 2015).

However, cereal production is significantly challenged by fungal diseases, which can cause substantial yield losses and reduce grain quality. Among the most concerning fungal pathogens are *Fusarium* spp., *Puccinia* spp. (rusts), and *Blumeria graminis* (powdery mildew), which not only compromise plant health but also lead to the accumulation of mycotoxins in grains, posing risks to human and animal health (Leonard and Bushnell, 2005; Różewicz et al., 2021; Moonjely et al., 2023). The economic impact of fungal diseases in cereals is substantial, requiring effective management strategies to minimize yield losses and ensure grain quality. The use of fungicides remains one of the most effective control measures, with strobilurins, azoles, and SDHIs (succinate dehydrogenase inhibitors) being commonly applied to suppress fungal growth (Poole and Arnaudin, 2014; Steinberg and Gurr, 2020; Jørgensen and Heick, 2021). Additionally, breeding for disease-resistant cultivars has become a key strategy, enabling crops to better withstand pathogen pressure and reducing the need for chemical interventions (Mesterhazy, 1995; Miedaner, 1997). An integrated approach combining resistant varieties, optimized fungicide applications, and agronomic practices is essential for maintaining sustainable productivity with high cereal yields while minimizing the risks associated with fungal diseases (Barzman et al., 2015).

Fungi of the genus *Trichoderma* are widely distributed in soil ecosystems and have long been recognized for their beneficial roles in agriculture (Druzhinina et al., 2010; Cai and Druzhinina, 2021). Several *Trichoderma* species are used as biocontrol agents due to their ability to suppress fungal plant pathogens through direct antagonism, competition for nutrients, and the production of antifungal metabolites (Harman, 2005). Commercial *Trichoderma*-based products are extensively applied as biologicals in crop protection, particularly against soilborne pathogens such as *Fusarium, Rhizoctonia*, and *Pythium*. In addition, some biologicals promote plant growth by enhancing root development and inducing systemic resistance (Sharma et al., 2012; Xue et al., 2017; Modrzewska et al., 2022). These characteristics have led to the widespread use of *Trichoderma* as an alternative to chemical fungicides considered environmentally friendly.

However, recent studies have revealed that certain *Trichoderma* species can also act as plant pathogen on maize, leading to severe symptoms on the cob and yield losses (Pfordt et al., 2020; Pfordt et al., 2025a). Trichoderma ear rot infected maize plants exhibit symptoms of white mycelium that rapidly produces dark, blue-green spores on maize cobs resulting in enzymatic decomposition of the kernel starch, making it soft and watery through the secretion of alpha-amylase (Pfordt et al., 2024). A monitoring conducted between 2018 and 2023 across Europe confirmed the widespread presence of *T. afroharzianum*, particularly in regions with warm and dry conditions in summer. The highest incidence of infected maize cobs and corresponding soil isolates was found in Southern Germany, France (Alsace), northeastern Austria, northern Italy and Türkiye. *T. afroharzianum* was consistently recovered from adjacent soils in affected regions, indicating the presence of the pathogen in agricultural soils. This raises important questions regarding the potential for cross-crop pathogenicity, especially in cereal-based crop rotations (Pfordt et al., 2025b).

Preliminary findings from greenhouse experiments indicate that *T. afroharzianum* is not strictly host-specific to maize. Under controlled conditions, the fungus was able to infect wheat and barley, leading to visible brown lesions on ears and kernels, as well as a measurable reduction in thousand kernel weight (Pfordt et al., 2023). These observations suggest that small grain cereals may represent additional hosts under conducive environmental conditions, particularly in warm and drought-prone regions.

The primary objective of this study is to investigate the pathogenicity of *T. afroharzianum* in different cereal crops under field conditions in order to validate and confirm previous greenhouse experiments which have demonstrated its ability to infect wheat and barley. To this end, the present study aims to evaluate the susceptibility of different wheat, barley, rye and triticale cultivar to *T. afroharzianum* infection and assess its effects on thousand kernel weight and colonization rates. Differences in host susceptibility among various cereal species and cultivars and the effectiveness of different fungicide treatments will be assessed.

By identifying differences in cultivar resistance and evaluating fungicide efficacy, this study aims to provide insights into effective disease management strategies to enhance the understanding and risk assessment of *T. afroharzianum* as a novel pathogen in cereal production systems. The demonstrated pathogenicity of *T. afroharzianum* challenges the long-standing perception of *Trichoderma* species as strictly beneficial organisms and underscores the necessity for a more differentiated perspective on their ecological roles and interactions with various host plants.

## 2. Materials and Methods

### 2.1 Used isolates and production of spore suspension

Three pathogenic *T. afroharzianum* isolates, obtained from maize were individually cultured on potato dextrose agar (PDA) for two weeks (Tab 1). Spores were harvested from PDA plates using a sterile spatula and suspended in sterile water. The spore concentration of each isolate was determined with a Thoma chamber and adjusted to 10^6^ spores/mL. Finally, the three spore suspensions were combined in equal proportions to generate one mixed inoculum (TriMix).

**Table 1.** Origin and year of isolation of the used *T. afroharzianum* isolates.

| Isolat | Species | Origin | Year |
| --- | --- | --- | --- |
| AP18TRI1 | *T. afroharzianum* | Maize cob, Croix des Pardis, France | 2018 |
| AP18TRI2 | *T. afroharzianum* | Maize cob, Kuenzing, Germany | 2018 |
| AP18TRI3 | *T. afroharzianum* | Maize cob, Pocking, Germany | 2018 |

### 2.2 Plant cultivation and experimental design

Two field experiments were conducted in 2023 and 2024 with a split-plot design, where fungicide treatment and varieties were the main plot factor, and inoculation (*Trichoderma* vs. water) was the subplot factor. Each cereal cultivar was assigned to a strip consisting of three adjacent plots, each receiving a different fungicide treatment. Within each plot, one half was designated for either *Trichoderma* inoculation or water treatment, serving as the subplot treatment.

The field trials were conducted at the Weendelsbreite site in Göttingen (51°33’47.0”N 9°56’51.9”E). The soil preparation involved conventional tillage, including plowing, followed by the use of a front-star roller and rotary harrowing to ensure a fine seedbed for optimal seed germination. Each experimental plot had an area of 12.5 m^2^ (2.5 m x 5 m). Fertilization was applied at a rate of 180 kg nitrogen per hectare (N/ha) across all plots to ensure adequate nutrient availability for crop growth throughout the trial period. The sowing of wheat, triticale, and rye in 2022 took place on October 5^th^, with barley sown between September 30th and October 3^rd^, 2022. In 2023, sowing for all the cereal species took place on October 4^th^ (Tab 2).

**Table 2.** Overview of sowing and flowering date in 2022/23 und 2023/24 of the used cereal varieties.

| Cereals | Cultivar | Breeder | Sowing<br>2022 | Flowering<br>2023 | Sowing<br>2023 | Flowering<br>2024 | Grain/<br>ha |
| --- | --- | --- | --- | --- | --- | --- | --- |
| Barley | Orbit | KWS | 30.09.22 | 16.05.23 | 04.10.23 | 14.05.24 | 300 |
|  | Higgins | KWS | 30.09.22 | 16.05.23 | 04.10.23 | 14.05.24 | 300 |
|  | Infinity | KWS | 02.10.22 | 22.05.23 | 04.10.23 | 15.05.24 | 300 |
|  | Quadriga | BayWa | 03.10.22 | 22.05.23 | 04.10.23 | 14.05.24 | 300 |
|  | Galileo | Syngenta | 01.10.22 | 22.05.23 | 04.10.23 | 14.05.24 | 300 |
| Rye | Dukato | SaatenUnion | 05.10.22 | 30.05.23 | - |  | 300 |
|  | Tayo | KWS | - |  | 04.10.23 | 23.05.24 | 300 |
| Triticale | Cosinus | KWS | 05.10.22 | 30.05.23 | 30.05.23 | 23.05.24 | 300 |
| Wheat | Asory | SECOBRA | 05.10.22 | 05.06.23 | 04.10.23 | 30.05.24 | 300 |
|  | Benchmark | Oberlimpurg | 05.10.22 | 05.06.23 | 04.10.23 | 28.05.24 | 300 |
|  | Chevignon | Hauptsaaen | 05.10.22 | 05.06.23 | 04.10.23 | 28.05.24 | 300 |
|  | Donovan | KWS | 05.10.22 | 10.06.23 | 04.10.23 | 04.06.24 | 300 |
|  | Euclide | Syngenta | 05.10.22 | 30.05.23 | 04.10.23 | 23.05.24 | 300 |
|  | Initial | Limagrain | 05.10.22 | 09.06.23 | 04.10.23 | 04.06.24 | 300 |
|  | Monopol | Firlbeck | 05.10.22 | 05.06.23 | 04.10.23 | 30.05.24 | 300 |
|  | Reform | Ragt | 05.10.22 | 08.05.23 | 04.10.23 | 28.05.24 | 300 |
|  | Ritmo | Limagrain | 05.10.22 | 09.06.23 | 04.10.23 | 04.06.24 | 300 |
|  | Tobak | Saaten Union | 05.10.22 | 09.06.23 | 04.10.23 | 04.06.24 | 300 |

### 2.3 Inoculation procedure

Inoculation of cereal heads was performed individually at full anthesis using a 5 L Gloria pump sprayer to ensure thorough coverage of cereal heads until runoff. Experimental plots were separated, with one side treated with the TriMix spore suspension and the other side sprayed with tap water as a control. To promote infection, ten to fifteen individual ears were grouped into bundles, with six to seven bundles per treatment. Immediately after inoculation, each bundle was enclosed in a polyethylene bag to maintain a humid microclimate conducive to fungal establishment. Bags were removed two days later.

### 2.4 Fungicide treatment

All plots were treated at the BBCH 09 growth stage (first leaf stage), with herbicide treatment to provide pre-emergence weed control before the winter season to prevent the growth of grassy and broadleaf weeds during the early stages of crop development. In plots 4 and 6, 0.75 kg/ha

‘Prodax’ was applied at the BBCH 31 growth stage (first node detectable). ‘Prodax’ contains trinexapac-ethyl and prohexadione-calcium, two growth regulators that inhibit gibberellin biosynthesis, leading to reduced stem elongation and increased stem stability (Tab. 3).

**Table 3.** Overview of herbicide und fungicide treatments applied to the plots during vegetation in 2023 and 2024.

| 2023 | Herbicide<br>12.10.2022. ES 09 | Growth promoter<br>18.04.2023. ES 31 | Fungicide 1<br>18.04.2023. ES31 | Fungicide 2<br>25.05.2023. ES 51 | Fungicide 3<br>05.06.2023. ES61 |
| --- | --- | --- | --- | --- | --- |
| **Plot 2** | 0.5 l/ha Cadou SC<br>0.35 l/ha Mateno Duo |  |  |  |  |
| **Plot 4** | 0.5 l/ha Cadou SC<br>0.35 l/ha Mateno Duo | 0.75 kg/ha Prodax | 1 l/ha Amistar | 1 l/ha Amistar |  |
| **Plot 6** | 0.5 l/ha Cadou SC<br>0.35 l/ha Mateno Duo | 0.75 kg/ha Prodax | 1.0 l/ha Input | 1.5 l/ha Ascra X Pro | 1.0 l/ha Input |
| 2024 | 10.10.2023. ES 09 | 09.04.2024. ES 31 | 09.04.2024. ES 31 | 21.05.2024. ES 51 | 27.05.2024. ES 61 |
| **Plot 2** | 0.6 l/ha Battle Delta |  |  |  |  |
| **Plot 4** | 0.6 l/ha Battle Delta | 0.4 l/ha Moddus<br>0.4 l/ha Medax Top | 1 l/ha Amistar | 1 l/ha Amistar |  |
| **Plot 6** | 0.6 l/ha Battle Delta | 0.4 l/ha Moddus<br>0.4 l/ha Medax Top | 1.0 l/ha Input | 1.5 l/ha Ascra X Pro | 1.0 l/ha Input |

\*ES Developmental stage

**Table 4.** Mean temperature (°C) and sum of precipitation (mm) from October until August in 2022/2023 and 2023/2024 in Goettingen, Germany.

|  |  |  | Oct | Nov | Dez | Jan | Feb | Mar | April | May | June | July | Aug |
| --- | --- | --- | --- | --- | --- | --- | --- | --- | --- | --- | --- | --- | --- |
| 2022/<br>2023 | Temp |  | 12.6 | 6.9 | 1.7 | 4.3 | 3.7 | 6.0 | 7.7 | 12.4 | 17.6 | 18.1 | 18.2 |
|  | Preci |  | 46.4 | 39.6 | 44.1 | 56.4 | 42.0 | 73.6 | 49.2 | 27.2 | 83.6 | 71.6 | 95.6 |
| 2023/<br>2024 | Temp |  | 12.5 | 6.1 | 4.8 | 1.9 | 7.2 | 7.4 | 10.1 | 15.1 | 16.1 | 18.4 | 19.8 |
|  | Preci |  | 89.2 | 77.6 | 134.4 | 62.0 | 70.0 | 44.0 | 60.0 | 64.8 | 70.4 | 118.0 | 92.4 |

In Plot 4, ‘Amistar’ (1 L/ha) was applied at first node stage and the beginning of ear emergence (BBCH 31 and BBCH 51). ‘Amistar’ contains azoxystrobin, a strobilurin fungicide effective against a broad spectrum of fungal pathogens, including rusts, powdery mildew, and *Fusarium* spp. It is primarily a protective fungicide with systemic properties. In Plot 6, ‘Input’ (1 L/ha) was applied at the first node stage (BBCH 31), followed by ‘Ascra X Pro’ (1.0 L/ha) at ear emergence (BBCH 51) and during flowering (BBCH 61/65). ‘Input’ is a fungicide that combines spiroxamine and prothioconazole, offering broad-spectrum control against fungal diseases such as *Septoria, Fusarium*, and rusts. It is a systemic fungicide with both protective and curative properties. ‘Ascra X Pro’ is a combination fungicide containing bixafen, prothioconazole, and fluopyram. ‘Ascra X Pro’ has systemic and translaminar movement activity in the plant, providing both preventive and curative action, and is particularly effective during critical growth stages such as flowering.

### 2.5 Sampling and harvest

Three weeks after inoculation, symptoms such as browning were observed in Plot 2, indicating infection. However, no systematic assessment of disease incidence or severity was conducted at this stage, as the observed symptoms could easily be confused with other diseases and were therefore not quantified. At maturity, 30 ears per treatment were selected and harvested in four replicates. The ears were cut, threshed, and then dried to a consistent moisture content.

### 2.6 Assessment of thousand kernel weight and colonization rate

Thousand kernel weight (TKW) was determined by weighing all harvested kernels from each plot and counting the total number of kernels. The total weight was then extrapolated to 1,000 kernels. To assess the colonization rate of *T. afroharzianum*, 100 kernels per cultivar and replication were randomly selected and surface disinfected with 0.25% silver nitrate for ten minutes. The kernels were then placed on folded sterile filter paper in incubation chambers and incubated in climate chambers at 25°C for five to seven days under sterile conditions. Filter paper remained moist by adding sterile water when necessary. After the incubation period, kernels showing fungal growth with characteristic green sporulation were counted. The colonization rate was calculated as the percentage of infected kernels in relation to the total number of analyzed kernels.

### 2.7 Statistical analysis

Data collected from the field trials were analyzed using TIBCO® Statistica software. The experiment was conducted using a split-plot design, with fungicide treatment as the main plot factor, crop cultivar as a blocking factor, and inoculation (*Trichoderma* vs. water) as the subplot factor. A linear mixed-effects model (LMM) was used to analyze the data, including the fixed effects of fungicide treatment, crop cultivar, and inoculation as well as the interactions between these factors. Random effects accounted for the plot structure and any variability due to blocks. Analysis of variance (ANOVA) between means of colonization rate and thousand kernel weight was carried out by Tukey-HSD-test at 5% probability. Statistical significance was determined at p < 0.05. Post-hoc pairwise comparisons were conducted using Tukey’s Honestly Significant Difference (HSD) test to evaluate differences between treatments. Data normality was assessed using the ShapiroWilk test. and homogeneity of variances was tested with Levene’s test. Results are presented as means ± standard errors (SE).

## 3. Results

### 3.1 Weather conditions

Temperature and precipitation data were evaluated for the periods October 2022 to August 2023 and October 2023 to August 2024, representing the respective growing seasons of winter cereals in Germany (Tab. 1).

In 2023, the average temperatures in May and June were 12.4 °C and 17.6 °C, respectively, accompanied by relatively low precipitation in May (27.2 mm) and moderate rainfall in June (83.6 mm). In contrast, 2024 showed a warmer pattern, with average temperatures of 15.1 °C in May and 16.1 °C in June. Precipitation was also higher in May 2024 (64.8 mm) but slightly lower in June (70.4 mm) compared to the previous year.

In 2022/2023, temperatures were near long-term averages in autumn and winter, with slightly warmer and wetter conditions in spring and summer. Overall, precipitation remained within typical annual ranges, while autumn and winter were somewhat cooler and drier than average. In contrast, 2023/2024 showed consistently higher temperatures across all seasons, with winter and spring exceeding long-term means by up to 3–4 °C. Summer 2024 was distinctly warmer, reaching 19.8 °C, and experienced significantly above-average precipitation, especially in July and August. Overall, 2023/2024 was markedly warmer and wetter compared to both the previous year and historical averages.

### 3.2 Disease symptoms

Three weeks after inoculation, first symptoms of *Trichoderma* infection were detected on some wheat cultivars. These included browning and light discoloration of the ears, particularly at the tips, as well as slight shriveling of some grains. As ripening progressed, the symptoms became increasingly difficult to distinguish from natural senescence. No symptoms were observed on barley, rye, or triticale. Other fungal infections, especially leaf diseases like *Septoria tritici, Drechselera triticirepentis*, and rust (*Puccinia* spp.), were also observed, varying in severity depending on the cultivar and fungicide application.

### 3.3. Effect of fungicide treatment on *Trichoderma* infection

Fungicide treatments exerted a significant effect (p < 0.0001) on the colonization rate and thousand kernel weight. All factors, such as fungicide treatment, inoculation and their interaction, were significant (p < 0.0001), whereas there was no significant effect of the replicate and year (Suppl.Tab.1). Results show that *Trichoderma* can infect small grain cereals under field conditions and reduce the TKW by increasing kernel colonization, especially in the absence of fungicidal treatment. As shown in Figure 1, the colonization rate of *Trichoderma* increased significantly in comparison to water-inoculated control plants, from 4% to 22%, which corresponded with a significant reduction of TKW by 11%, from 35 g to 31 g.

**Figure 1.**
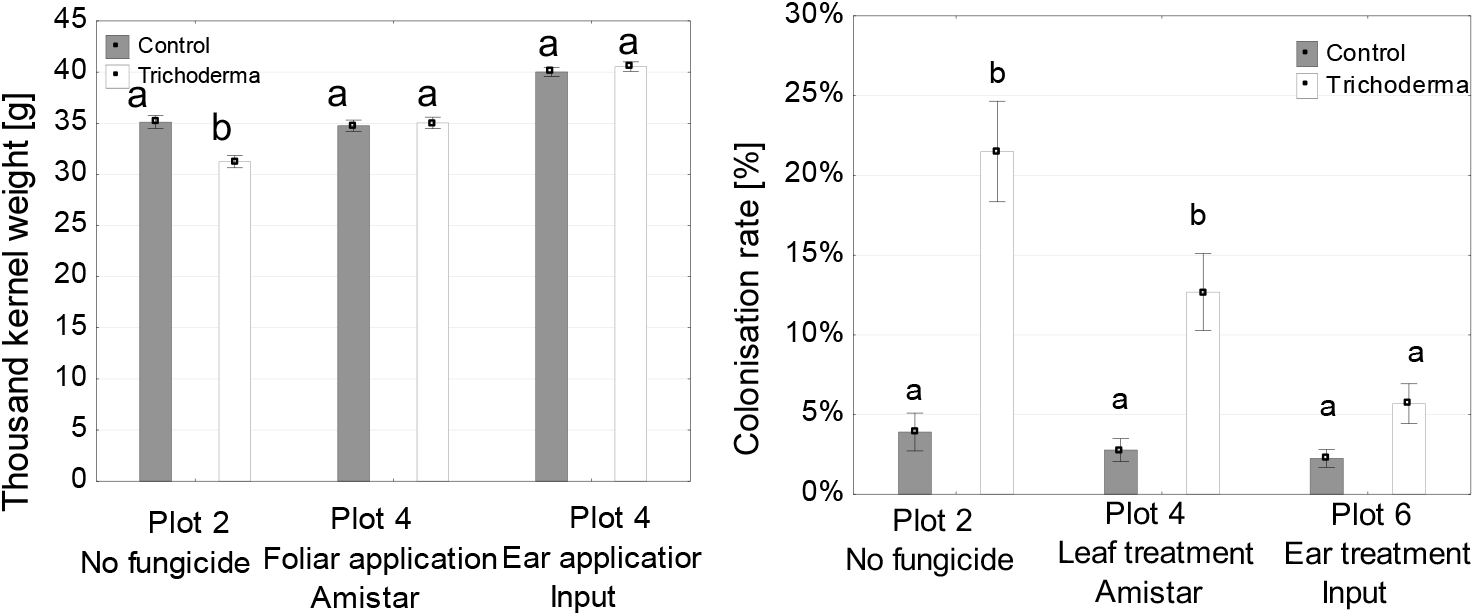
Thousand kernel weight (g, left) and colonization rate (%, right) on all grains at different fungicide treatments in Plot 2 (no fungicide treatment), Plot 4 (leaf treatment with ‘Amistar’ at BBCH 31 and BBCH 51), and Plot 6 (fungicide treatments with Input at BBCH 31, ‘Ascra X Pro’ at BBCH 51, and ‘Input’ at BBCH 61) after inoculation with *T. afroharzianum* and water as control. Diffferent letters indicate significant differences between treatments across plots. Bars represent standard errors, asterisks represent significant differences between treatments (α=0.05, Tukey HSD test).

In the foliar treatment (Plot 4), colonization rates also increased significantly from 3% to 13% after application of ‘Amistar’ at growth stages BBCH 31 and BBCH 51. However, this increase did not result in a reduction of TKW. In the ear treatment (Plot 6), which received three fungicide applications with ‘Input’ (BBCH 31), ‘Ascra XPro’ (BBCH 51), and ‘Input’ during flowering (BBCH 61), neither colonization rates nor TKW showed significant changes compared to the control. In general, ear application with ‘Input’ increased the TKW compared to both the untreated control and the foliar treatments, likely by effectively reducing infection from other fungal diseases.

### 3.4 Susceptibility of cereal species to *Trichoderma* infection

The results indicate that cereal species and inoculation treatments as well as their interaction had a significant (p < 0.03375) impact on TKW and colonization rate (Suppl. Tab 1).

Figure 2 illustrates the impact of *T. afroharzianum* infection on TKW and colonization rate across different cereal species after icoulation with *T. afroharzianum* und water (control). The most notable decrease is found in wheat where the infected grains exhibit a significantly lower TKW (-6 g) compared to water inoculated control plants. Slight reductions in TKW are observed for barley, while rye and triticale showed no significant differences. The highest colonization rates were observed in wheat and barley with infection rates exceeding 10% and peaking at around 17% in barley. Rye and triticale exhibited much lower colonization rates, suggesting that these species may be more resistant to *T. afroharzianum* colonization.

**Figure 2.**
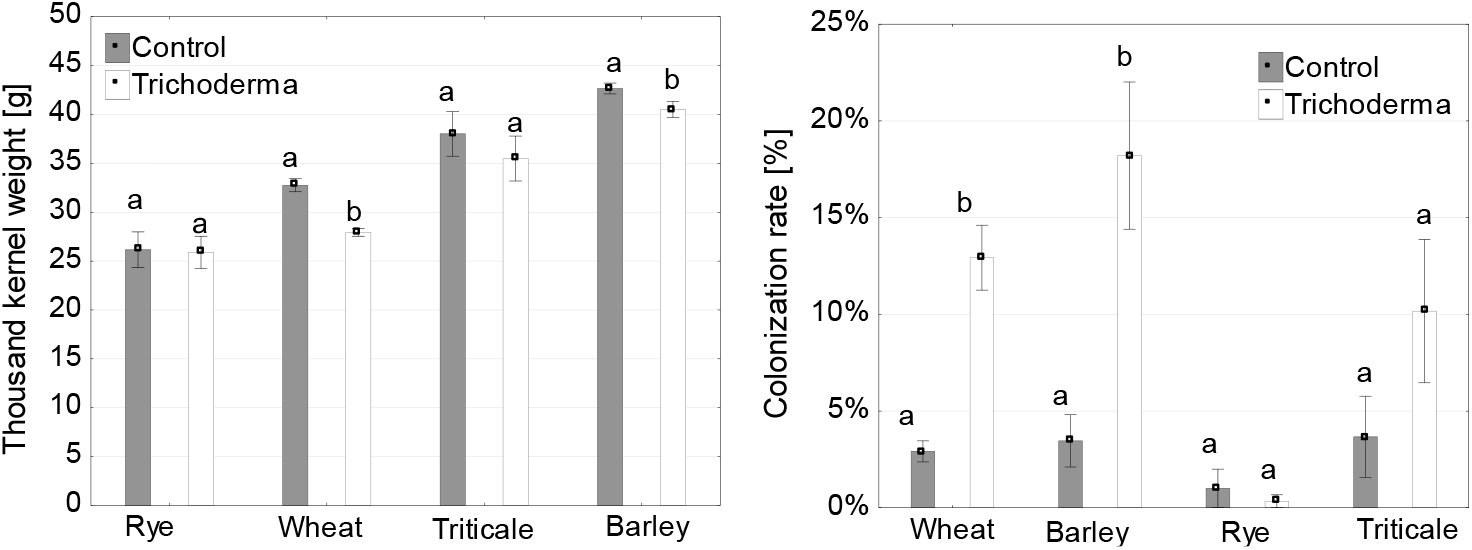
Thousend kernel weight (left) and % colonisation rate (right) on grains of rye, wheat, barley and triticale after incoulation with *T. afroharzianum* and water as control without fungicide treatment (plot 2). Bars represent standard errors, asterisks represent significant differences between treatments (α=0.05, Tukey HSD test).

### 3.5 Cultivar effects on *Trichoderma* infection

The thousand kernel weight of different cereal varieties, including barley, rye, triticale, and wheat differed significantly (p < 0.0001), following inoculation with *T. afroharzianum* and treatment with water (control)(Figure 3). Among the barley cultivars, only ‘Galileoo’ exhibited a significant decrease in TKW after *Trichoderma* infection. The remaining barley cultivars, ‘Orbit’, ‘Higgin’, and ‘Quadriga’, did not show significant changes compared to the control. Both rye cultivars, ‘Toya’ and ‘Dukato’, showed no noticeable reduction in TKW under *Trichoderma* treatment. The triticale cultivars ‘Cosinus’ displayed a moderate response, with slight reductions in TKW that were not statistically significant. Wheat cultivars showed more variable responses. Most cultivars, including ‘Benchmark’, ‘Donovon’, ‘Reform’, ‘Chevignon’, ‘Tobak’, ‘Monopol’, and ‘Infinity’, experienced significant reductions in TKW, whereas ‘Initial’, ‘Ritmo’, ‘Euclinde’, and ‘Asory’ were unaffected by the infection.

**Figure 3.**
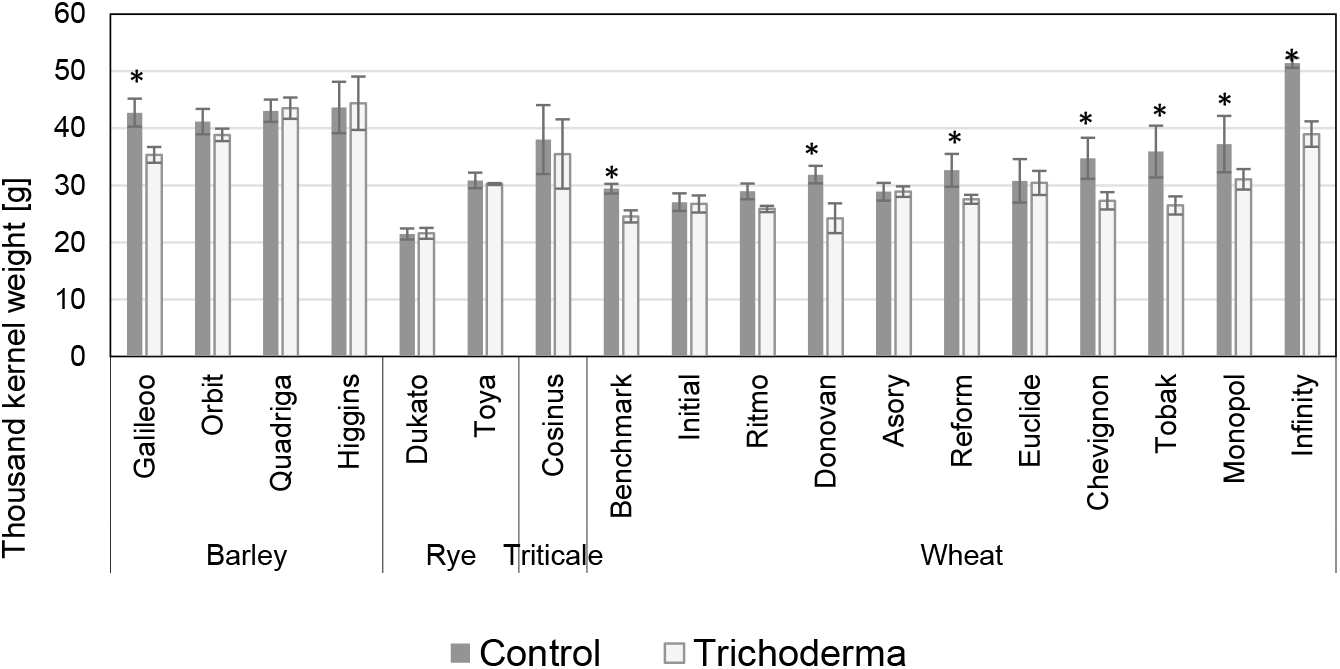
Thousand kernel weight of barley, rye, triticale and wheat cultivars after inoculation with *T. afroharzianum* and treatment with water as control and. Bars represent standard errors, asterisk represent significant differences between the treatments (α=0.05, Tukey HSD test).

Figure 4 illustrates the colonization rates (%) of *T. afroharzianum* across different cereal varieties following inoculation with *Trichoderma*. Among the barley varieties, ‘Galileo’ showed the highest colonization rate, reaching nearly 50%. In contrast, ‘Orbit’ and ‘Quadriga’ exhibited lower, yet still significant, levels of fungal colonization, while ‘Higgins’ appeared largely resistant, with no noticeable increase in infection compared to the control. Both rye varieties showed no increase in colonization rates, and the triticale cultivar displayed a slight increase relative to the control. Wheat varieties demonstrated more variable responses. ‘Tobak’ exhibited the highest colonization rate, exceeding 40%, while ‘Ritmo’, ‘Initial’, ‘Reform’, ‘Asory’, and ‘Monopol’ showed moderate to high increases in colonization compared to the control.

**Figure 4.**
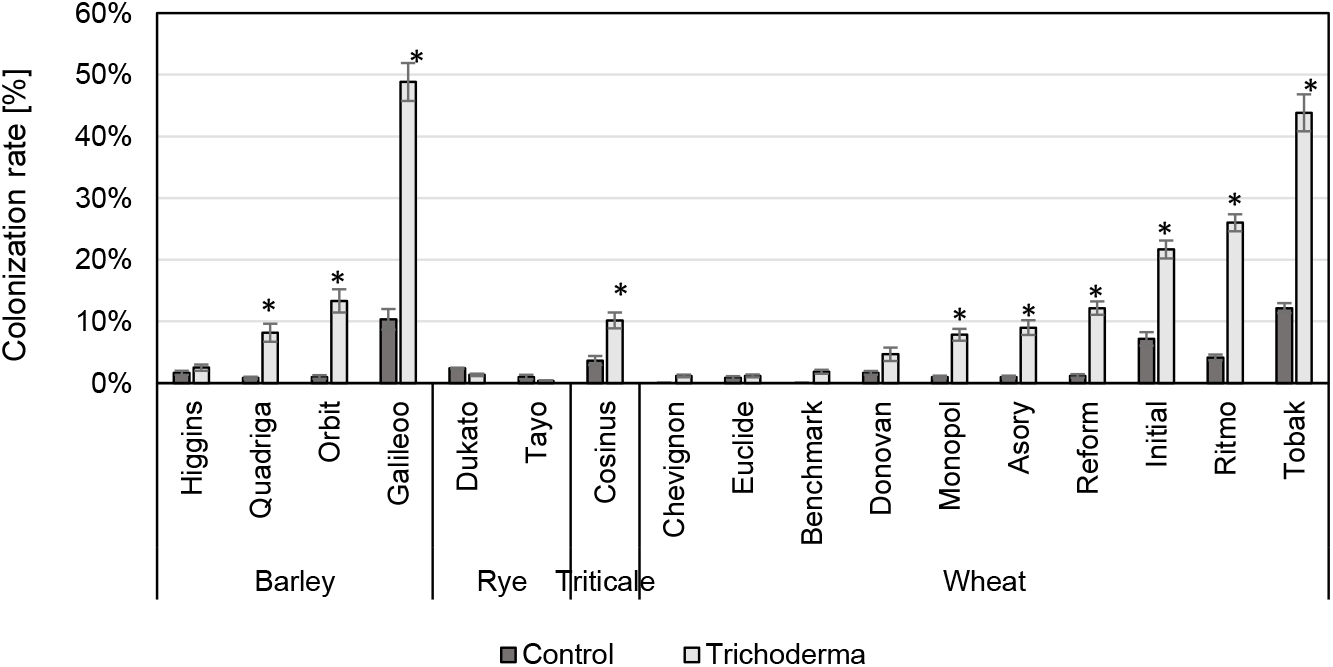
Colonization rate (%) on grains of barley, rye, triticale and wheat varieties after inoculation with *T. afroharzianum* and treatment with water as control and. Bars represent standard errors, asterisk represent significant differences between treatments (α=0.05, Tukey HSD test).

## 4. Discussion

The present study provides the first field-based evidence that *T. afroharzianum*, previously characterized mainly as a biocontrol agent, can act as a pathogenic fungus on small grain cereals under field conditions. During two years of field trials, several wheat and barley cultivars exhibited significant reductions in thousand kernel weight and displayed high colonization rates, while rye and triticale remained largely unaffected. Furthermore, fungicide treatments revealed differential effectiveness in mitigating infection, with ear-targeted applications being most successful. These findings extend previous greenhouse observations by demonstrating that *T. afroharzianum* is capable of colonizing multiple cereal species under field conditions, although the extent of damage is strongly influenced by host genotype, fungicide use, and environmental factors.

The findings indicate that certain rye and wheat varieties are particularly susceptible to *T. afroharzianum*, whereas barley and triticale exhibit lower infection rates. The results align with the TKW data, showing that varieties with a significant decrease in TKW also exhibit a high colonization rate. This suggests a direct relationship between fungal infection intensity and yield reduction. For example, barley cultivar ‘Galileo’ and wheat varieties ‘Tobak’ and ‘Ritmo’ display both high colonization rates and notable reductions in TKW, indicating that *T. afroharzianum* significantly impacts grain development in these varieties. In contrast, rye and triticale varieties, which showed lower colonization rates, maintained relatively stable TKW, suggesting a higher level of resistance or tolerance to the pathogen.

The effectiveness of fungicide treatments against *T. afroharzianum* infection strongly depends on the timing and mode of action of the applied fungicides. The differences observed between the plots suggest that fungicide application before infection primarily influenced fungal colonization, while post-infection treatments had a greater impact on both colonization and grain quality. In plot 2, where no fungicide was applied, *T. afroharzianum* colonization reached 22%, which was significantly higher than in the fungicide-treated plots. This high colonization rate coincided with a significant reduction in thousand kernel weight (TKW), confirming the strong negative effect of *T. afroharzianum* infection on grain development. These results indicate that, in the absence of fungicidal protection, the pathogen can successfully colonize cereal ears and negatively affect yield formation. Plot 4, which received ‘Amistar’ (azoxystrobin) as a foliar treatment before infection, showed a colonization rate of 12%, demonstrating that the treatment did not fully prevent fungal spread. However, TKW was not significantly reduced, suggesting that despite the increased colonization, the overall grain development remained unaffected. Azoxystrobin, a strobilurin fungicide, primarily acts as a protective agent by inhibiting mitochondrial respiration in fungal cells, thereby preventing spore germination and early infection stages. However, because it was applied before flowering and infection it likely had no direct effect on *T. afroharzianum* colonization within the ears. The results indicate that while the treatment may have suppressed early pathogen growth, it was not sufficient to fully prevent ear colonization. In contrast, Plot 6, which received ‘Ascra Xpro’ (bixafen + fluopyram + prothioconazole) as an ear treatment after infection, exhibited the lowest colonization rate (6%) and prevented any significant TKW reduction. The effectiveness of this treatment can be attributed to its systemic and curative properties. Prothioconazole, a demethylation inhibitor (DMI) fungicide, disrupts ergosterol biosynthesis, inhibiting fungal cell membrane formation and stopping pathogen development within plant tissues. Bixafen and fluopyram, both succinate dehydrogenase inhibitors (SDHIs), block mitochondrial respiration, thereby limiting fungal energy production. This combination provides both protective and curative effects, reducing colonization even after infection has already occurred.

The effectiveness of fungicide treatments observed in this study against *T. afroharzianum* shows striking parallels to findings from Fusarium head blight (FHB) research. For FHB, numerous field trials and meta-analyses have demonstrated that fungicide efficacy is highly dependent on application timing, with the greatest reductions in disease severity and mycotoxin contamination achieved when sprays are applied at early anthesis and up to a few days thereafter (Jones, 2000). Triazole fungicides such as prothioconazole, tebuconazole, and metconazole have consistently provided the strongest activity against FHB, and their effectiveness is further enhanced in resistant cultivars (Paul et al., 2019). More recently, co-formulations of DMIs with SDHIs, such as bixafen + fluopyram + prothioconazole (‘Ascra Xpro’), have been shown to perform equally well or better under field conditions (Edwards, 2022). In contrast, strobilurins such as azoxystrobin are regarded as unreliable for FHB control and in some cases have been associated with unfavorable effects on mycotoxin accumulation (Haidukowski et al., 2005). This suggests that future disease management strategies should prioritize fungicides with systemic activity and curative properties, particularly for pathogens that infect cereal ears during flowering (Mesterházy et al., 2011; D’Angelo et al., 2014). The results highlight key differences in the effectiveness of fungicide treatments against *T. afroharzianum*, with significant implications for practical agriculture. In conventional farming, fungicide applications targeting both leaf and head diseases, particularly against *Fusarium* spp., are standard practice (Bernhoft et al., 2022). These treatments also effectively reduce *Trichoderma* colonization and minimize yield losses. Consequently, organic farming, where chemical fungicides are not permitted during flowering, faces a higher risk of *Trichoderma* infections (van Bruggen et al., 2016). Increased colonization rates and associated yield reductions highlight the need for alternative disease management strategies in organic systems. The development of resistant varieties and improved agronomic practices, such as optimized crop rotation and soil management, will be essential to mitigate *Trichoderma*-related losses in cereal production.

In the present studies, wheat and barley were susceptible to *Trichoderma* colonization, whereas rye and triticale remained largely unaffected. A similar pattern is known from other ear diseases, most prominently Fusarium head blight were wheat and barley are typically the most susceptible species, while rye and triticale generally show lower levels of infection. These parallels suggest that host specificity may be shaped by similar factors across different pathogens, including floral traits, ear architecture, and intrinsic plant defense responses. In both *Trichoderma* and *Fusarium* spp., dense and moisture-retaining ears of wheat and barley promote colonization, whereas the looser ear structure and stronger basal resistance of rye and triticale restrict pathogen establishment.

The results indicate a potential correlation between the susceptibility of wheat cultivars to *Fusarium* and their colonization by *T. afroharzianum*. According to assessments by the German Federal Plant Variety Office (Bundessortenamt), the wheat varieties ‘Ritmo’ and ‘Tobak’ are classified as highly susceptible to Fusarium head blight (rating 7), whereas all other tested wheat varieties have moderate susceptibility ratings (4 or 5). This classification aligns with our findings, as ‘Tobak’ (44%), ‘Ritmo’ (26%) and ‘Initial’ (22%) exhibited the highest *T. afroharzianum* colonization rates, while varieties with lower *Fusarium* susceptibility scores, such as ‘Benchmark’ (2%), ‘Chevignon’ (1%), and ‘Euclide’ (1%), showed minimal colonization. This suggests that wheat varieties with weak resistance to *Fusarium* may also be more vulnerable to *T. afroharzianum* infection and that there may be cross resistance to the two ear pathogens. This is in contrast to another ear disease, wheat blast caused by *Magnaporthe oryzae triticum* pathotype, where no such positive cross resistance with fusarium head blight was found (Ha et al., 2016).

Future research should focus on monitoring naturally *Trichoderma*-infected wheat in organic production systems to assess the prevalence and impact of the pathogen under non-chemical management conditions. This will provide insights into whether *T. afroharzianum* may pose a recurring threat in organic farming and how environmental factors influence its spread and severity. Additionally, investigating host resistance mechanisms across different cereal species and varieties is crucial. Understanding genetic and physiological traits that contribute to resistance are essential to support breeding efforts for more resilient cultivars. These research directions will help develop effective management strategies and improve the sustainability of both organic and conventional cereal production.

The results highlight that *Trichoderma afroharzianum* is a potential novel cereal ear pathogen which may significantly impact cereal production, particularly in susceptible varieties and under conditions where no effective fungicide treatments are applied. While rye and triticale appear relatively tolerant, wheat and some barley cultivars showed greater susceptibility, suggesting that host resistance varies across cereal species and cultivars. Fungicide applications, particularly those targeting *Fusarium* during flowering, effectively also reduced *Trichoderma* colonization and protected grain quality in a conventional farming system. In organic production, where fungicide treatments are not available, *Trichoderma* could become a more significant problem, necessitating alternative disease management strategies such as resistant varieties, optimized crop rotations, and biological control practices. These findings underscore the importance of integrating host resistance, fungicide treatments, and agronomic practices to safeguard cereal yields and quality against emerging fungal threats.

## Author Contributions

methodology. S. PB.; validation. A.P.; investigation. A.P.; writing—original draft preparation. A.P.; writing—review and editing. A.v.T.; visualization. A.P.; supervision. A.v.T.; project administration. A.P.; funding acquisition. A.P and A.v.T. All authors have read and agreed to the published version of the manuscript.”

## Ethics approval and consent to participate

Not applicable

## Consent for publication

Written informed consent has been obtained from the authors to publish this paper

## Availability of data and material

The datasets used and/or analyzed during the current study are available from the corresponding author on reasonable request.

## Funding

This research was funded by Federal Ministry of Agriculture, Food and Regional Identity, grant number FKZ2221NR014

### Acknowledgments

We would like to thank Manuela Mücke for technical support of the experiments and for performing DNA sequencing, respectively.

## Supplemental Material

**Suppl. Table 1.** ANOVA of fungicide treatment, cultivar, *Trichoderma* inoculation, cereals, year and replication on colonization rate and thousand kernel weight.

|  | Treatment | df | F | PR>F |
| --- | --- | --- | --- | --- |
| Colonisationrate | Fungicide treatment | 2 | 20.62 | < 0.0001 |
|  | Cultivar | 14 | 11.94 | < 0.0001 |
|  | *Trichoderma* inoculation | 1 | 55.71 | < 0.0001 |
|  | Cereal | 3 | 2.910 | 0.03375 |
|  | Year | 1 | 3.985 | 0.12843 |
|  | Replication | 3 | 0.812 | 0.48721 |
| Thousand-kernel-weight | Fungicide treatment | 2 | 168.0 | < 0.0001 |
|  | Cultivar | 14 | 17.92 | < 0.0001 |
|  | *Trichoderma* inoculation | 1 | 7.066 | 0.00801 |
|  | Cereal | 3 | 12.84 | 0.03845 |
|  | Year | 1 | 0.119 | 0.73017 |
|  | Replication | 3 | 1.246 | 1.69147 |

